# *In vitro* and *in vivo* derived macrophages occupy distinct phenotypic states

**DOI:** 10.1101/2025.09.20.677495

**Authors:** Stéphane Chevrier, Vito R. T. Zanotelli, Daniel Schulz, Mark D. Robinson, Laurie Ailles, Michel A.S. Jewett, Craig Gedye, Bernhard Reis, Bernd Bodenmiller

**Author notes:** Correspondence should be addressed to B.B.

## Abstract

Monocyte-derived macrophages (MDMs) are widely used to model human macrophage biology *in vitro* and standardized polarization conditions were proposed to recapitulate the different macrophage activation states. Although surface markers specific for distinct MDM populations have been identified, a systematic analysis of surface marker expression on these consensus MDM populations has not previously been reported. Here, we use mass cytometry to perform an in-depth characterization of MDM surface profiles and determine how these markers evolve with time. We also compared the phenotypes found *in vitro* with patient-derived tumor-associated macrophages (TAMs) and found that although MDMs and TAMs shared most markers investigated, the cell-surface signatures markedly differed in terms of expression levels and combinations. Our high-dimensional, single-cell analyses clarifies the surface expression profile of the core *in vitro* differentiated macrophage populations and highlights some limitations of the *in vitro* system to represent the complexity of *in vivo* polarized tumor-associated macrophage phenotypes. These findings provide new opportunities to improve the models used to study macrophage biology and give a pathway to improve our understanding of the *in vivo* complexity of these systems.

## INTRODUCTION

Macrophages are distributed throughout the body and are involved in a wide variety of tasks including clearance of senescent cells, angiogenesis, bone resorption, elimination of allergens, clearance of pathogens, antigen presentation, inflammation, and tissue repair ^1–3^. Abnormal macrophage functions can lead to or reinforce pathological conditions, as observed in metabolic diseases, allergic disorders, autoimmune diseases, and cancer ^1,2^. Macrophages have been often segregated into classically activated M1 and alternatively activated M2 subtypes, which are assumed to constitute two extremes of a continuum ^4–6^. M1 macrophages are characterized by the secretion of pro-inflammatory cytokines, reactive oxygen species, and nitric oxide, which mediate their microbicidal and tumoricidal activities. M2-like macrophages secrete immunosuppressive cytokines as well as growth and angiogenic factors and are proposed to be involved in tissue remodeling and resolution of the inflammation ^7^.

*In vitro* models using monocyte-derived macrophages (MDMs) have been developed to mimic the functional phenotypes found *in vivo*. The polarization stimuli used to generate MDMs have evolved from the original M1/M2 dichotomous model induced using IFNγ and IL4, respectively, to include glucocorticoid, immune complexes, and additional cytokines. These differentiation conditions resulted in better defined subsets such as M2a, M2b, and M2c ^8,9^. The growing numbers of MDM populations have been extensively studied at the transcriptional and protein levels using microarrays, deep sequenced transcriptome, proteomics as well as multiplexed cytokine/chemokine secretion assays ^10–14^. These studies reported that macrophage polarizations cover a spectrum of activation states ^5,15^. To avoid confusion in the terminology used and to allow better comparison of experimental data across laboratories and disciplines, standardized *in vitro* polarization conditions and nomenclatures were proposed ^16^. The core polarizations that cover the spectrum of macrophage activation states were defined, and the authors proposed that these activation states could serve as a framework to map *ex vivo* isolated macrophages ^16^. Cell surface marker signatures are a key tool for mapping and analyzing macrophage types from diseased tissues within a clinical context. Although previous studies have identified surface markers specific for distinct MDM populations, no systematic analyses have been performed to characterize and catalogue the surface expression profile of those core MDM activation states, thus hindering their further assessment and utility.

Unlike most immune cell types, macrophage subsets cannot be defined based on the expression or the lack of expression of a few well-defined markers ^16–18^. Due to their inherent plasticity, a large number of markers are required to distinguish the phenotypic spaces occupied by macrophage subsets. Mass cytometry, which makes use of metal-tagged antibodies, enables the simultaneous detection of up to 40 protein readouts at the single-cell level with limited signal overlap ^19–21^. Moreover, this method is not affected by the auto-fluorescence of myeloid cells, which can lead to artifacts in classic fluorescence flow cytometry. With the ability to measure dozens of samples and millions of cells in a short time, mass cytometry is a method-of-choice to directly profile and compare the surface expression profiles of differentially activated macrophages.

To systematically analyze the surface expression profiles of MDMs generated under standard conditions and in a time resolved manner ^16^, we employed an adapted version of a 36-antibody panel used to characterize tumor associated macrophage in the context of renal carcinoma^22^ in a mass cytometry analysis. We then directly compared the surface phenotypes of MDMs with the phenotypes of tumor-associated macrophages (TAMs) from ascites isolated from ovarian cancer patients. Although the surface markers identified by analysis of MDMs were detected on TAMs, we observed little overlap in the actual surface marker expression patterns. Our data highlight the complex phenotypic diversity of macrophages found *in vivo*, emphasizing the limitations of the *in vitro* MDM model to recapitulate *in vivo* macrophage states.

## RESULTS

### MDM characterization using mass cytometry

In order to develop an antibody panel that best discriminate blood monocytes (M0), immature macrophages (M5), and populations obtained by stimulation with the nine MDM conditions proposed by Murray and colleagues ^16^ (Figure 1A), we selected markers based on a fluorescent surface screening previously performed on those populations ^22^. The final antibody panel (Table 1) was used to analyze simultaneously the expression of 36 markers on the surface of eleven populations of monocytes and monocyte derived macrophages from four healthy donors. In order to minimize technical variability during staining and to measure all samples simultaneously, the samples were barcoded using a 60-sample mass-tag cell barcoding method with a doublet filtering scheme ^23,24^.

**Table 1.** Surface molecules included in the final panel.

| Metal Tag | Target | Antibody Clone | Biological function |
| --- | --- | --- | --- |
| La139 | CD209 | DCS-8C1 | C-type lectin involved in phagocytosis |
| Pr141 | CD64 | 10.1 | High affinity Fc receptor |
| Nd142 | CD38 | HIT2 | cyclic ADP ribose hydrolase |
| Nd143 | CD68 | Y1/82A | glycoprotein |
| Nd144 | CD36 | 5-271 | scavenger receptor class B |
| Nd145 | CD71 | CY1G4 | Transferrin receptor I |
| Nd146 | CD155 (PVR) | SKII.4 | Costimulatory receptor |
| Sm147 | CD206 | 15-2 | Mannose receptor |
| Nd148 | CD13 | WM15 | Alanine aminopeptidase |
| Sm149 | CD204 | 351615 | Scavenger receptor |
| Nd150 | CD123 | 6H6 | Interleukin-3 receptor |
| Eu151 | CD11b | M1/70 | Complement receptor 3 |
| Sm152 | CD40 | 5c3 | Costimulatory receptor |
| Eu153 | Slamf7 | 162.1 | Costimulatory ligand |
| Sm154 | CD197 | GO43H7 | Chemokine receptor |
| Gd155 | CD273 | MIH18 | coinhibitory receptor |
| Gd156 | CD304 | 446921 | VEGF co-receptor |
| Gd158 | CD166 | 3A6 | Adhesion molecule |
| Tb159 | CD82 | ASL-24 | Tetraspanin glycoprotein |
| Gd160 | CD274 | 130021 | coinhibitory ligand |
| Dy161 | CD119 | GIR-208 | Interferon-γ receptor |
| Dy162 | CXCR4 | 12G5 | Chemokine receptor |
| Dy163 | CD87 | VIM5 | Urokinase receptor |
| Dy164 | CD120b | 3G7A02 | Tumor necrosis factor receptor 1B |
| Ho165 | CD4 | RPA-T4 | T cell coreceptor |
| Er166 | CD32 | FUN-2 | Low affinity Fc receptor |
| Er167 | CD16 | 3G8 | Low affinity Fc receptor |
| Er168 | CD14 | RMO52 | LPS co-receptor |
| Tm169 | CD163 | GHI/61 | Scavenger receptor |
| Er170 | CD169 | 7-239 | Syaloadhesin |
| Yb171 | CD86 | 2331 (FUN-1) | Costimulatory ligand |
| Yb172 | HLA-ABC | W6/32 | Antigen presentation |
| Yb173 | CD81 | 5A6 | Tetraspanin glycoprotein |
| Yb174 | CD88 | S5/1 | complement component 5a receptor 1 |
| Lu175 | HLA-DR | L243 | Antigen presentation |
| Yb176 | CD54 | HA58 | Adhesion molecule |

**Figure 1.**
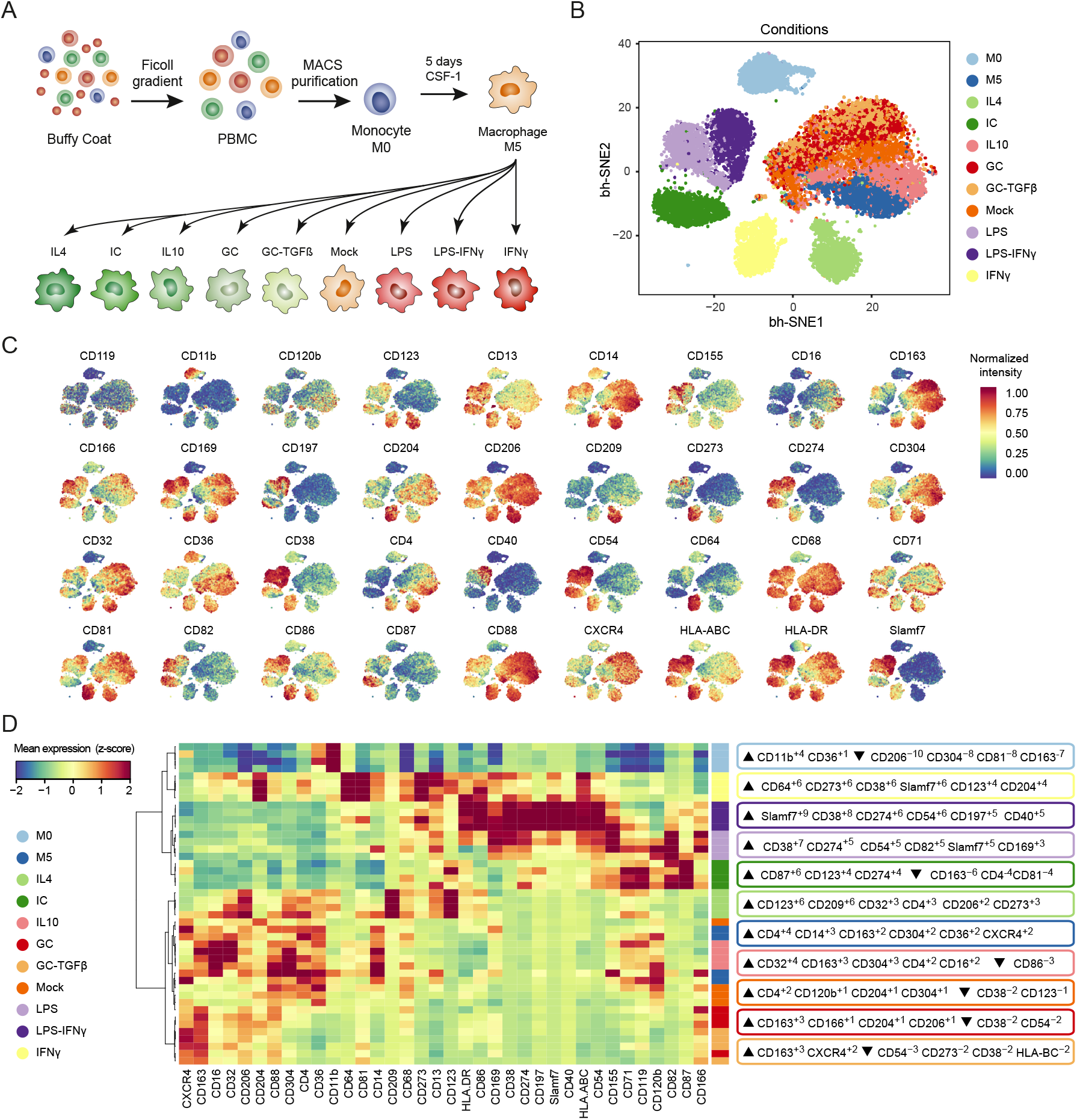
Characterization of MDM surface phenotypes by mass cytometry. (A) Monocyte preparation and stimulation to induce differentiation. (B) t-SNE map of M0 and MDM populations based on the 36 antibodies used for mass cytometry analysis. Cells are colored by sample types. (C) t-SNE map showing the normalized expression levels of the 36 surface markers across the different populations. Data are from one representative replicate.(D) Heatmap showing the normalized expression level of the 36 antibodies for the four replicates of the different populations included in the analysis. Hierarchical clustering was performed by calculating first the pairwise distance between samples based on the Spearman correlation, and by clustering based on Ward’s method. For each subset, the six proteins showing the strongest enrichment based on marker enrichment modelling is indicated on the right. IC, immune complex; GC, glucocorticoid. See also Figure S1 and Table S1.

Using the nonlinear dimensionality reduction approach based on the t-distributed stochastic neighbor embedding (t-SNE) ^25,26^, the high-dimensional single-cell data were visualized in two dimensions (Figure 1B). This approach showed that monocytes, M(IL4), M(Immune complex (IC)), and M(IFNγ) were clearly separated from all other populations. M(LPS) and M(LPS-IFNγ), although distinct, were localized in close proximity. M5, M(Mock), M(IL10), M(Glucocorticoid (GC)), and M(GC-TGFβ) showed partial overlap. This configuration was consistent for each of the four donors (Figure S1A).

Unique patterns for each marker assessed in this study were observed in the t-SNE map (Figure 1C). The canonical macrophage markers CD68, CD14, and CD13 were expressed in all populations, yet at different abundances. CD11b was high only in monocytes and downregulated upon *in vitro* culture. As expected from the original optimization of the antibody panel, we identified markers that were population specific, such as CD209 in M(IL4), CD87 in M(IC), and CD64 in M(IFNγ). Most markers were shared across a range of conditions, however; and this was particularly obvious for costimulatory molecules. CD40 was induced both in M(LPS) and M(LPS-IFNγ) culture conditions. CD155, Slamf7, and CD274 (PD-L1) were expressed in the M1 subsets (LPS, LPS-IFNγ, and IFNγ) and in the M(IC). CD273 (PD-L2) and CD86 were induced in M1 and M(IC) populations and showed some expression in the M(IL4). Similarly, the adhesion molecules CD54 (ICAM-1) and CD82, the cyclic ADP ribose hydrolase CD38, and the chemokine receptor CCR7 were also strongly induced in the M1 and the M(IC) subsets. The IL3 receptor CD123 was expressed on both M1 and M(IC) populations and was observed at the highest level in M(IL4). The scavenger receptors CD163 and CD36, the chemokine receptor CXCR4, the Fc receptor CD32, the VEGF co-receptor CD304, and the complement receptor CD88 had strong expression in the M2-like populations polarized with CSF-1, IL4, IL10, GC, and GC-TGFβ. The CD4 activation marker had an expression pattern overlapping with typical M1 and M2 subsets and showed a high induction upon IL4, IFNγ, and IL10 treatment. The data were consistent across the four samples analyzed (Figure S1B). Despite the heterogeneity of monocytes used to generate macrophages, all the markers used in the analysis showed a unimodal (albeit sometimes broad) distribution on all the MDM populations investigated (Figure S1C).

Hierarchical clustering showed that despite the biological variation, most MDM donor replicates clustered based on the activator used. To systematically identify the marker signature that best characterizes the different MDM populations, we used marker enrichment modelling, a metric that takes into account the median intensity and the distribution of the markers in each population ^27^. The marker signature identified for each subset based on this approach was largely consistent with the observation made based on the t-SNE map, as shown in Figure 1D. Taken together, our data showed that the antibody panel used captures MDM heterogeneity and establishes condition-specific surface expression signatures, which can be used to compare samples across studies.

### Evolution of MDM surface marker expression over time

To understand better the dynamic nature of marker expression over time, we harvested MDMs 6h, 12h, 24h, 36h and 48h after activation with the different stimuli. Displaying the cells from the different time points for each stimulation on a t-SNE map revealed that the *in vitro* differentiation process is dynamic and phenotypic changes could be observed throughout the entire kinetic (Figure 2A, Figure S2A). A systematic analysis of the log2 fold change of each marker at the different time points upon stimulations compared to the initial state (M5) revealed that most markers were induced upon stimulation whereas only a minority of markers were repressed (Figure 2B). For most markers, the induction or the repression was reinforced throughout the kinetic and the highest or lowest level was reached after 48h. The markers that best defined the M5 signature (Figure 1D), including CD4, CD14, CD163, but also CD32 or CD206 were among the markers that were repressed upon stimulation (Figure 2B). The repression of those markers was occurring mostly on the M1 populations (LPS, LPS-IFNγ, and IFNγ) and the M(IC) while their expression levels were maintained or even slightly increased on the M2-like subsets. This was consistent with the fact that the CSF-1 cytokine used to generate the M5, triggers an anti-inflammatory M2-like response and therefore many M2 specific markers were already induced on M5 cells. For most markers induced during the kinetic, including CD64, CD81, CD38, CD40, CD197, CD54, CD274, CD273, CD166 or CD82, the expression level showed a progressive increase over at least the first 24h of activation. One notable exception was Slamf7, which was already expressed at its maximum level only 6h after LPS or LPS-IFNγ stimulation and 12h after IC activation (Figure 2B).

**Figure 2.**
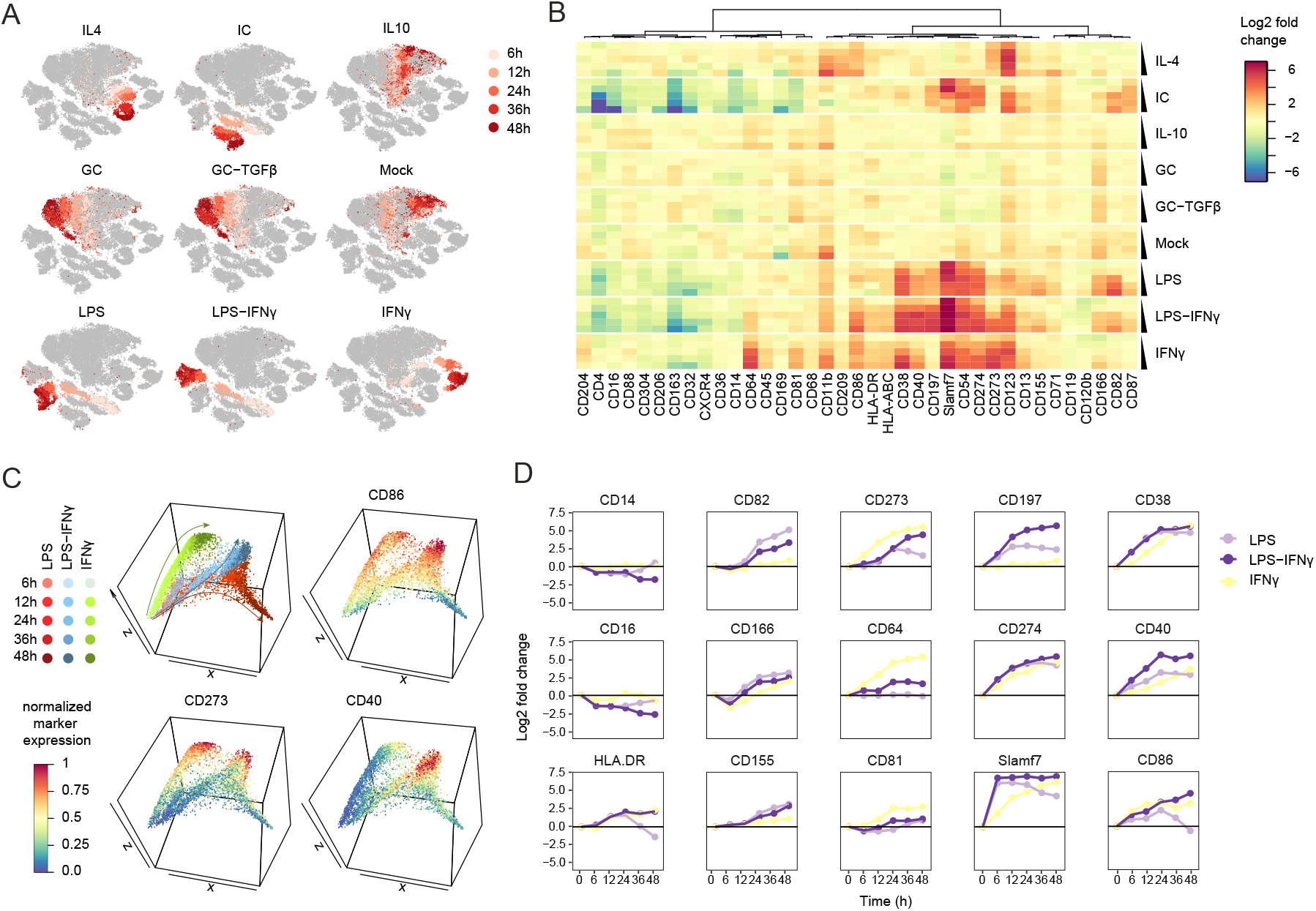
Kinetic analysis of MDM polarization. (A) t-SNE maps of MDMs activated with the indicated stimuli for the indicated amount of time. For each stimulation, the cells are colored with a gradient of red corresponding to the vstimulation time shown on a grey background corresponding to all the other cells included in the analysis. (B) Heatmap showing the log2 fold change in marker expression across time points and stimuli compared to M5. Markers are clustered based on Spearman correlation and Ward’s linkage. (C) Diffusion mapping based on the first three diffusion components showing the putative trajectories followed by monocytes activated with the indicated stimuli over 48h (top-left panel). The other panels are showing the diffusion map colored by the expression level of three markers differentially expressed on the different trajectories. (D) Line plot showing the log2 fold change of the indicated marker in monocyte population activated with LPS, LPS-IFNγ and IFNγ for the indicated time points compared to M5.

When looking specifically at cells activated with LPS, LPS-IFNγ and IFNγ alone, both the t-SNE and the fold change analysis suggested strong similarities between cells activated with LPS and with LPS-IFNγ during the first 12h (Figure 2A-B). A further analysis based on diffusion mapping, an algorithm that aligns cells along putative developmental trajectories confirmed that monocytes activated with LPS and LPS-IFNγ followed the same trajectory during the first time points before adopting different differentiation paths in the later time points (Figure 2C). Conversely, cells activated with IFNγ followed a different trajectory from the beginning (Figure 2C). This suggested a dominant role of LPS at earlier time points, while the effect of IFNγ in presence of LPS was mostly observed after 24h. The presence of IFNγ was critical to induce a high expression level of costimulatory molecules including CD86, CD273 and CD40 and to maintain a long lasting induction of Slamf7 on LPS treated cells (Figure 2C). When looking systematically at the evolution of marker expression at the different time points, we observed that different canonical macrophage markers, including CD14, CD16 or HLA-DR were indeed expressed at the same levels on cells activated with LPS and LPS-IFNγ for up to 24h before diverging at the later time points (Figure 2D, first column). This approach also revealed different regulatory mechanisms for the different markers investigated. LPS was a more potent inducer of CD82, CD166 and CD155 than IFNγ or the combination of both (Figure 2D, second column). On the other hand, IFNγ had a stronger effect on the induction of CD273, CD64 or CD81 (Figure 2D, third column) compared to LPS alone or in combination with IFNγ. Finally, for most of the costimulatory molecules, including CD274, CD40, SLAMF7, CD86, but also for the ecto-enzyme CD38 and the chemokine receptor CCR7 (CD197), the combination of both LPS and IFNγ was required for a sustained, high expression level (Figure 2D, fourth and fifth columns). Together, these analyses enabled to get a better understanding on the constant evolution of marker expression over time and demonstrated how LSP, IFNγ and the combination thereof differentially regulated the expression of key macrophage markers.

### MDMs and ascites associated macrophages have distinct phenotypes

The transcriptional and phenotypic signatures of different MDM populations were used to map myeloid cells isolated from human pathological conditions ^5,28^. However, understanding to which extend MDM populations can serve as a framework to map and classify macrophages isolated from patient samples is still an open question. We used our comprehensive MDM dataset to directly assess how close *in vitro* derived macrophages resemble to *in vivo* isolated tumor associated macrophages. In order to avoid tissue dissociation, a step susceptible to modify the native phenotype found in vivo, we focused this comparison on ascites from advanced ovarian cancer patients, known to contain high amount of macrophages. Since ascites were stored as viably frozen cells, we compared the profile of MDMs before and after freezing and we observed no significant changes (Figure S2 A-B).

We analyzed the phenotypes of macrophages isolated from ascites of nine ovarian patients simultaneously with the MDM populations. To allow the detection of T cells and macrophages, the panel was complemented with antibodies to CD3 and CD45. TAMs were identified as CD45^+^ CD3^-^ CD68^+^ CD64^+^ live cells and T cells as CD45^+^ CD3^+^ (Figure S2C). The phenotypes of MDMs, TAMs, and T cells were compared based on a t-SNE analysis on those populations. This approach showed that a small proportion of macrophages had an antibody staining signature similar to that of M0 (blood monocytes) consistent with the fact that tumors are vascularized and likely contain circulating monocytes (Figure 3A). However, when we compared the signatures of ascites associated macrophages with those of the MDMs, no overlap was observed. This indicated that macrophages differentiated *in vitro* do not recapitulate the cell-surface protein phenotypes of macrophages found in the ascites samples (Figure 3A).

**Figure 3.**
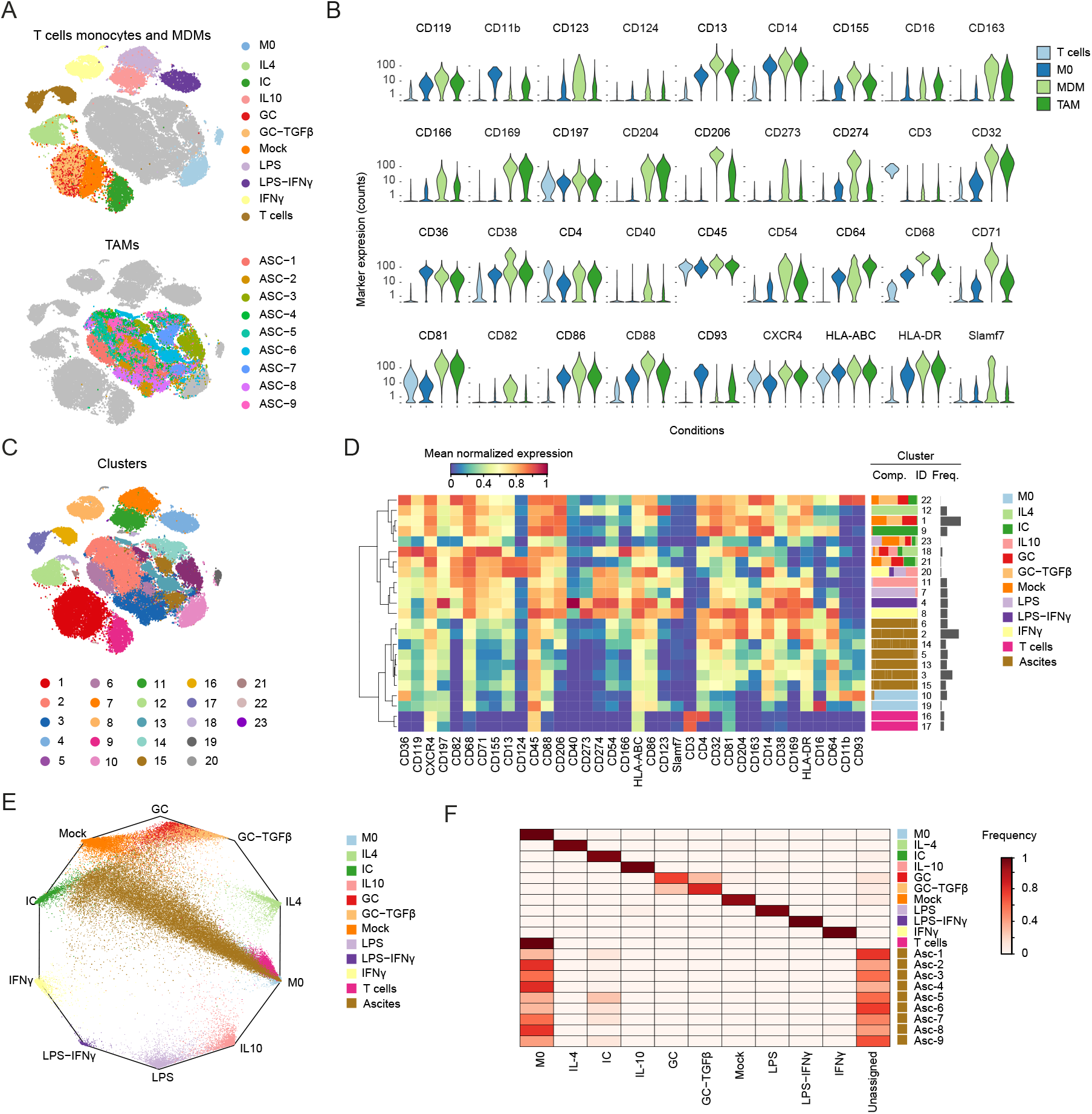
MDMs and TAMs express different combinations of similar markers. (A) t-SNE maps of TAMs, T cells, M0, and MDM populations based on the CyTOF antibody panel. Cells were colored per MDM sample types (top panel) or per ascites samples (bottom panel). In each case, the counterpart populations are colored in grey. (B) Violin plot showing the expression pattern of the 36 markers across T cells, monocytes, pooled MDMs, and pooled TAMs. For pooled populations, data on 2,000 cells of each individual sample were combined. (C) t-SNE map of the 36 markers across T cells, monocytes, pooled MDMs, and pooled TAMs colored by PhenoGraph clusters. (D) Heatmap showing the mean expression levels of the 36 markers assessed by mass cytometry across the 26 PhenoGraph clusters. Three clusters with less than 100 cells were left out of the analysis. Hierarchical clustering was performed by calculating first the pairwise distance between samples based on the Spearman correlation and by clustering based on Ward’s method. The composition (comp.) and the frequency (freq.) of each cluster are depicted on the right. (E) Wheel plot showing the prediction score of MDMs, TAMs and T cells, based on a random forest classifier trained on the MDM populations. The distance between each dot, representing a cell and the vertices of the polygons is proportional to the predication score of each cell to the different MDM populations. (F) Heatmap showing for each cell type the frequency of cells classified with a 40% confidence into the different population used to train the model. See also Figure S2.

The absence of overlap between TAMs and MDMs on the t-SNE map could be explained if the markers chosen based on *in vitro* differentiated macrophages were not expressed on the surface of TAMs. We therefore compared the expression levels of all the markers used in our study, on the surface of T cells, monocytes, MDMs, and macrophages. For this comparison, data on all the MDM populations were pooled as were data on macrophages from the nine ascites samples. As expected, T cells were positive only for CD4, CD81, CXCR4, and HLA-ABC (Figure 3B). The monocytes expressed high levels of CD11b, CD13, CD14, and CD36 and intermediate levels of CD68, CD64, and HLA-DR. The pooled MDM population showed detectable expression for all the markers investigated except CD11b. Strikingly, the expression pattern of the different markers on the surface of the TAM population obtained by pooling the nine samples was similar to the expression pattern observed for MDM populations. In particular, the expression patterns of CD166, CD32, CD38, CD81, CD204, HLA-ABC, and HLA-DR were almost indistinguishable between ex vivo isolated macrophages and MDMs. Some markers, such as CD4 and CD64 were expressed at higher levels on ascites associated macrophages than on MDMs. Although some markers, including CD206, CD163, and CD169, were expressed at lower levels on ex vivo isolated macrophages than on MDMs, the expression was still higher than on monocytes, and a proportion of cells expressed a level close to the level observed on MDMs. Costimulatory molecules (such as CD40, CD274, and Slamf7) and chemokine and cytokine receptors (such as CCR7 and CD119) were significantly reduced on ex vivo isolated macrophages compared to MDMs. These data show that the markers analyzed were appropriate for the characterization of *in vivo* macrophage phenotypes.

As an alternative approach to better understand the absence of surface marker phenotype overlap between TAMs and MDMs and to exclude the possibility that t-SNE leads to misleading conclusions, we identified ex vivo isolated macrophages and MDM subsets using the unsupervised graph-based clustering algorithm PhenoGraph ^30^. Consistent with t-SNE-based observations, the ex vivo isolated macrophages - and MDM-containing clusters were mutually exclusive, whereas one cluster contained both TAMs and monocytes (Figure 3D, middle panel, cluster 10). Ex vivo isolated macrophages subsets were characterized by subtle differences in a wide variety of surface markers with only a few markers reaching a high expression level. In contrast, the MDM subsets were characterized by clear differences in marker expression with multiple markers expressed at high levels. Ascites associated clusters were characterized by different combinations of markers that could not individually be associated with specific MDM populations.

To assess the similarity between ascites associated cells and *in vitro* derived macrophages, we developed a machine learning approach comparing the marker signature of the cells found *in vivo* with the one generated *in vitro* based on a random forest classifier. Using a wheel plot, we visualized each cell based on its probability of being assigned to each of the MDM populations used to train the model (Figure 2E). Apart from the cells activated with glucocorticoid alone or in presence of TGFB, which are difficult to distinguish, cells from each MDM subset were located in close vicinity to the correct node. This was further confirmed when looking at the frequency of cells assigned to the different cell types based on a 40% probability threshold (Figure 2F). In comparison, macrophages isolated from ascites formed a range of poorly defined cells, originating from the M0 monocyte subset, but rapidly losing any connections with *in vitro* derived cell subsets (Figure 2E). Consistently, most of the macrophages from ascites samples were assigned to the monocyte subset, or left unassigned based on a 40% probability threshold (Figure 3F). Thus, our data show that the markers identified based on MDM populations do capture ex vivo macrophage diversity but are expressed in different combinations and often at a lower level leading to an absence of phenotype overlap between ex vivo macrophages and MDM subsets.

## DISCUSSION

Macrophages are highly plastic cells that respond to stimuli present in the local environment by adopting a continuum of functional phenotypes ^3,31^. To comprehend this complexity in humans, numerous studies over the last two decades have relied on the MDM *in vitro* system and the M1/M2 conceptual framework (Mills et al., 2000; Martinez and Gordon, 2014). Although this system has been used to evaluate transcriptional networks and chemokine and cytokine profiles of MDMs, no studies have systematically investigated the surface expression profiles of MDM populations, a necessary step to determine how the *in vitro* system relates to macrophages *in vivo*.

We used a systematic approach to characterize the cell-surface phenotypes of the nine core MDM populations proposed by Murray and colleagues (Murray et al., 2014) based on a modified version of a recently developed mass cytometry antibody panel ^22^. Despite differences in culture protocols, our results were mostly consistent with previous reports, with features such as an increased expression of CD11b and CD206 on M(IL4) and an upregulation of CD64, Slamf7, and CD86 on M(IFNγ) as previously described ^10,33^. Our strategy extended the surface expression signature to a larger panel of markers and to more MDM populations and was consistent across different donors.

Since most markers included in the panel have well-established biological functions, this characterization furthered our understanding of the functional differences between the MDM subsets. In a time course analysis, we observed that LPS has a dominant effect on IFNγ during MDM activation and that cells activated with LPS alone or in presence of IFNγ displayed strongly overlapping phenotypes up to 12 hours after activation. At later time points, these two MDM subsets still share many features. Our approach also showed how LPS or IFNγ alone or in combination differentially controls the expression of costimulatory molecules such as CD40, Slamf7, CD273 and CD274, highlighting their complex regulation during macrophage activation. Although MDM activation by immune complexes was previously associated with an anti-inflammatory, M2-like phenotype ^16^, our data suggest that these cells share surface features with pro-inflammatory cells including induction of the costimulatory molecules CD155, CD274, and Slamf7 and expression of CD38, CD54, CD82, and CD197.

We then took advantage of the newly defined MDM signatures to determine whether MDM phenotypes could be used as a framework to classify *ex vivo* isolated TAM populations ^22^. We compared the cell-surface phenotypes of MDM populations with TAMs isolated from ccRCC tumors from 20 patients using the nonlinear dimensionality reduction algorithm t-SNE, which enables the visualization of high-dimensional single-cell data on a 2D map that groups cells based on their similarity, and the cluster algorithm PhenoGraph. This approach showed that MDM populations failed to reproduce the phenotypes of TAMs found *in vivo*, and our attempt to use machine learning to classify ascites associated macrophage into the different MDM subsets showed that ex vivo macrophages could not be classified into any of the populations derived *in vitro*.

Due to the strictly controlled culture conditions, involving the presence of only one or two cytokines, it is perhaps not surprising that *in vitro* populations did not mimic the phenotypes of macrophages found in the very complex *in vivo* environment where cells are simultaneously exposed to a multitude of stimulations and where various feedback mechanisms are at work. This likely explains the fact that many markers, especially costimulatory molecules and chemokine receptors, were induced at higher levels in MDMs compared to TAMs.

Comparing macrophages activated *in vitro* with macrophages isolated *ex vivo* is inherently limited by the fact that samples are treated in ways that can potentially prevent a direct comparison. To circumvent this issue, we focused here on macrophages isolated from ascites samples, which do not require any tissue dissociation, susceptible to affect marker expression due to the enzymatic treatment and to further stimulate macrophages due to the presence of dying cells. A limitation of our study is that we only analyzed surface markers, whose expression level is more likely to change compared to more hardwired markers, such as key transcription factors or even cytokine secretion. The detection of such markers is limited by antibody availability, the sensitivity of the system, and the need to reactivate the cells, all of which can influence results.

Although the MDM system has proven valuable in the study of macrophage biology over the past decades, our study highlights the limitation of this system. A better understanding of macrophages present in pathological conditions requires a deeper characterization based on direct analysis of *ex vivo* isolated cells. The advent of mass cytometry, which allows simultaneous analysis of multiple functional markers, has enabled the characterization of cell subsets on a substantial scale. Combining mass cytometry with other high-dimensional analyses, such as RNA sequencing or multiplexed cytokine assays, has already started to uncover biological features of macrophage subsets in complex tissues ^22,34^. More effort in this direction is critical to better understand the role of macrophages in various pathological conditions and to facilitate the development of new therapeutic treatments.

## MATERIALS AND METHODS

### Monocyte isolation and macrophage generation

Buffy coats from healthy donors were obtained from the Zurich Blood Transfusion Service. PBMCs isolation was performed by histopaque (Sigma Aldrich) density gradient centrifugation, and monocytes were isolated by MACS using the pan monocyte isolation kit (Miltenyi Biotech) according to manufacturer’s instructions. Monocytes (M0) were differentiated into immature macrophages (M5) by culture for 5 days in Nunc UpCell™ 6-well plates in presence of 30 ng/ml M-CSF (PeproTech). Cells were subsequently polarized for the indicated amount of time in fresh medium complemented with the following stimuli: 20 ng/ml IL4 (PeproTech), 10 ng/ml IL10 (PeproTech), 10 μg/ml IC (Sigma Aldrich), 40 ng/ml of the GC dexomethasone (Sigma Aldrich), 40 ng/ml GC (Sigma Aldrich) and 10 ng/ml TGFβ (CST), 100 ng/ml LPS (Sigma Aldrich), 100 ng/ml LPS (Sigma Aldrich) and 100 ng/ml IFNγ (PeproTech), and 100 mg/ml IFNγ (PeproTech). Mock-treated cells were cultured for the same amount of time with fresh culture medium only. After culture, cells were harvested by temperature reduction. For mass cytometry analysis, cells were stained for viability with 10 µM cisplatin (Enzo Life Sciences) in a 1-min pulse before quenching with 10% FBS as previously described ^35^. Cells were then fixed with 1.6% paraformaldehyde (Electron Microscopy Sciences) for 10 min at room temperature. Cells were then stored at −80 °C.

### Tumor sample preparation

Primary ascites samples from ovarian cancer patients were obtained from the xxx from consenting patients under University Health Network Research Ethics Board approval. Tumors were graded based on were histological analysis by a pathologist (reported in Table S3). Single-cell were collected by centrifugation, resuspended in D-SDCM and counted with a hemocytometer. Cells were then frozen in 10% DMSO in D-SDCM complemented with 10% FBS. Cryopreserved cells were resuscitated for mass cytometry analyses by rapid thawing and slow dilution. Cells were stained for viability with 10 µM cisplatin (Enzo Life Sciences) in a 1-min pulse before quenching with 10% FBS as previously described ^35^. Cells were then fixed with 1.6% paraformaldehyde (Electron Microscopy Sciences) for 10 min at room temperature. Cells were then stored at −80 °C.

### Fluorescence cell barcoding and antibody screening

Fixed cells were washed in 0.03% saponin in PBS and incubated in PBS containing 0, 0.01, 0.1, or 1 µg/ml Alexa Fluor 700 carboxylic acid, succinimidyl ester (Molecular Probes) and 0, 0.1, or 10 µg/ml of Alexa Fluor 750 carboxylic acid, succinimidyl ester (Molecular Probes) for 15 min on ice, enabling the barcoding of up to 16 samples ^37^. Cells were then washed twice with 0.03% saponin in PBS, and the samples were pooled. The combined samples were screened with the BioLegend human cell screening kit containing 342 pre-titrated PE-conjugated antibodies arrayed on four 96-well plates according to manufacturer’s instructions. To avoid nonspecific staining, cells were first incubated for 10 min at 4 °C with human Fc-receptor (FcR) blocking reagent (Miltenyi Biotech). Cells were washed, filtered, and resuspended at a density of 4.0 × 10^6^ cells/ml. Approximately 2 × 10^5^ cells were added to each well of the BioLegend plates, and samples were incubated for 20 min at room temperature for antibody staining. After staining, cells were washed and fixed. Immediately after staining, data were acquired on a Canto II (BD Biosciences) using the auto sampler. Data were analyzed in Cytobank 4.10 as previously described ^38^.

### Mass cytometry barcoding

To ensure homogenous staining, 0.5 × 10^6^ cells from each sample were barcoded prior to cell staining and mass cytometry analyses. For this purpose, cells were labeled with a unique combination of four of eight barcoding reagents, enabling the simultaneous analysis of up to 60 samples as previously described ^23^. Six palladium isotopes (^102^Pd, ^1042^Pd, ^105^Pd, ^106^Pd, ^108^Pd, and ^110^Pd, Fludigm) were conjugated to bromoacetamidobenzyl-EDTA (BABE) and two indium isotopes (^113^In and ^115^In, Fludigm) were conjugated to 1,4,7,10-tetraazacy-clododecane-1,4,7-tris–acetic acid 10-maleimide ethylacetamide (mDOTA) following standard procedures ^39^. Mass tag barcoding reagents were titrated to ensure an equivalent staining for each reagent; final concentrations were between 50 nM and 200 nM. Cells were barcoded using the transient partial permeabilization protocol described by Behbehani and colleagues ^40^. Briefly, approximately 0.5 × 10^6^ fixed cells were loaded into the appropriate number of wells in a 96-well plate. Cells were washed with 0.03% saponin in PBS and incubated for 30 min with 200 μl of mass tag barcoding reagent. After staining, cells were washed twice with 0.03% saponin in PBS and twice with cell staining medium (CSM, PBS with 0.5% bovine serum albumin and 0.02% sodium azide). Samples were then pooled for subsequent cell staining.

### Antibodies and antibody labeling

Provider, clone, and metal tag of each antibody used in this study are listed in Table S1. Antibodies were labeled with the indicated metal tag using the MaxPAR antibody conjugation kit (Fluidigm). After metal conjugation, the concentration of each antibody was assessed using the Nanodrop (Thermo Scientific) and adjusted to 200 μg/ml in Candor Antibody Stabilizer. Conjugated antibodies were titrated to determine the optimal concentration for use. All antibodies used in this study were managed using the cloud-based platform AirLab_41_.

### Antibody staining and data acquisition

After barcoding, pooled cells were incubated with FcR blocking reagent (Miltenyi Biotech) for 10 min at 4 °C. Samples were stained with 200 µl of the antibody panel per 10^7^ cells for 30 min at 4 °C. Cells were washed twice in CSM and resuspended in 1 ml of nucleic acid Ir-Intercalator (Fluidigm) in 1.6% PFA/PBS for 1 h at room temperature or overnight at 4 °C. Cells were then washed twice in PBS and twice in water. Cells were then diluted to 0.5 × 10^6^ cells/ml in water containing 10% of EQ™ Four Element Calibration Beads (Fluidigm). Data were acquired on a CyTOF 2 mass cytometer, using instrument-based dual-count calibration and noise reduction and randomization, and cells were selected based on event length between 10 and 75. Exported flow cytometry standard (FCS) files were concatenated using the FCS concatenation tool from Cytobank and normalized using the executable MATLAB version of the Normalizer tool ^42^. Individual samples were debarcoded using the executable MATLAB version of the single cell debarcoder tool ^23^.

## Data analysis

FCS files were uploaded into Cytobank, populations of interest were manually gated, and events of interest were exported as FCS files. For downstream analysis, FCS files were loaded into R ^43^. Signal intensities for each channel were arcsinh transformed with a cofactor of 5 (x_transf = asinh(x/5)). To visualize the high-dimensional data in two dimensions, the t-SNE algorithm was applied on data from 2000 random cells from each sample. The total cell population was used when less than 2000 cells were available. The R t-SNE package for Barnes-Hut implementation of t-SNE was used ^44^. Data were displayed using the ggplot2 R package ^45^. To visualize marker expression analyses on t-SNE map, the top percentile was excluded and the maximum intensity was defined as the 99^th^ percentile. The data from all samples were divided by this value leading to signal intensities ranging between 0 and 1 for each channel. For hierarchical clustering, pairwise distances between samples were calculated using the Spearman correlation. Dendrograms were generated using Ward’s method and arranged according to the ‘order.optimal’ function of the ‘cba’ R package using optimal leaf ordering ^46^. Heatmaps were displayed in R using the heatmap.2 function form the gplot package. For single-cell clustering analysis, the Python implementation of PhenoGraph was run on all samples simultaneously with the parameter *k* defining the number of nearest neighbors set to 30 ^30^. For single cell classification, a random forest classifier (RandomForest R package) was trained on 3,000 cells of each MDM subsets using the default parameters (500 trees and 6 variables tried at each split). The remaining cells of the MDM populations, as well as the total populations of macrophages and T cells found in the ascites samples were classified based on this model. Probability assignment of each cell was visualized on a wheel plot (Radviz R package).

## ACKNOWLEDGEMENTS

We are very grateful for the generous donation of tumor samples by patients undergoing surgery, collected through the University Health Network Biobank and Cooperative Health Tissue Network. The authors thank the Bodenmiller lab for fruitful discussions and the Cytof facility (University of Zurich). B.B.’s research is funded by a SNSF R’Equip grant, a SNSF Assistant Professorship grant, the SystemsX Transfer Project “Friends and Foes”, the SystemsX MetastasiX grant, and by the European Research Council (ERC) under the European Union’s Seventh Framework Program (FP/2007-2013)/ERC Grant Agreement n. 336921. S.C. was funded by a Roche Postdoctoral Fellowship; D.S. by an EMBO fellowship (ALTF 970-2014) co-funded by the European Commission (LTFCOFUND2013, GA-2013-609409); C.G. by an RACP CSL Fellowship, a CIHR/KCC SHOPP Fellowship, and an NHMRC Early Career Fellowship.

## AUTHOR CONTRIBUTIONS

S.C., B.R., and B.B. conceived the study. S.C. performed all experiments with help from D.S. L.A., M.A.S.J., and C.G. S.C., V.R.T.Z., M.D.R. performed data analysis. S.C., L.A., M.A.S.J., C.G., B.R., and B.B. performed the biological analysis and interpretation. S.C. and B.B. wrote the manuscript with input from all authors.

## FIGURE LEGENDS

**Figure S1. Comparison of Marker Expression across the Different MDM Populations, Related to Figure 1**

(A) The data on MDMs generated from monocytes from four healthy donors were analyzed simultaneously using the t-SNE statistic. For each sample, the data are displayed individually on a scatter plot. In each graph, the indicated replicate is displayed using the indicated color code, and the three other replicates are shown in grey. (B) Median expression levels of the indicated markers in monocytes and MDM populations across four replicates. (C) Violin plots showing the distribution of the indicated markers in the different MDM populations as assessed by mass cytometry. Data were normalized to the 99 percentile. Data shown are from one representative sample.

**Figure S2. Effect of Cell Freezing on Marker Expression as Assessed by Mass Cytometry, Related to Figure 2**

(A) Gating strategy based on Ir, Pt, CD45, CD3, CD64, and CD68 used to identify T cells and TAMs. (B) t-SNE maps of the different MDM populations based on the CyTOF antibody panel in presence (right panel) or in absence (left panel) of freezing step. Cells were colored per MDM sample types and in each case, the counterpart populations are colored in grey. (C) Correlation between marker expression in presence or in absence of enzymatic digestion in the indicated cell samples. For each marker, the coefficient of determination R2 and the linear model are displayed.

