## Supplementary material for "*In vitro* and *in vivo* derived macrophages occupy distinct phenotypic states": Figure S1

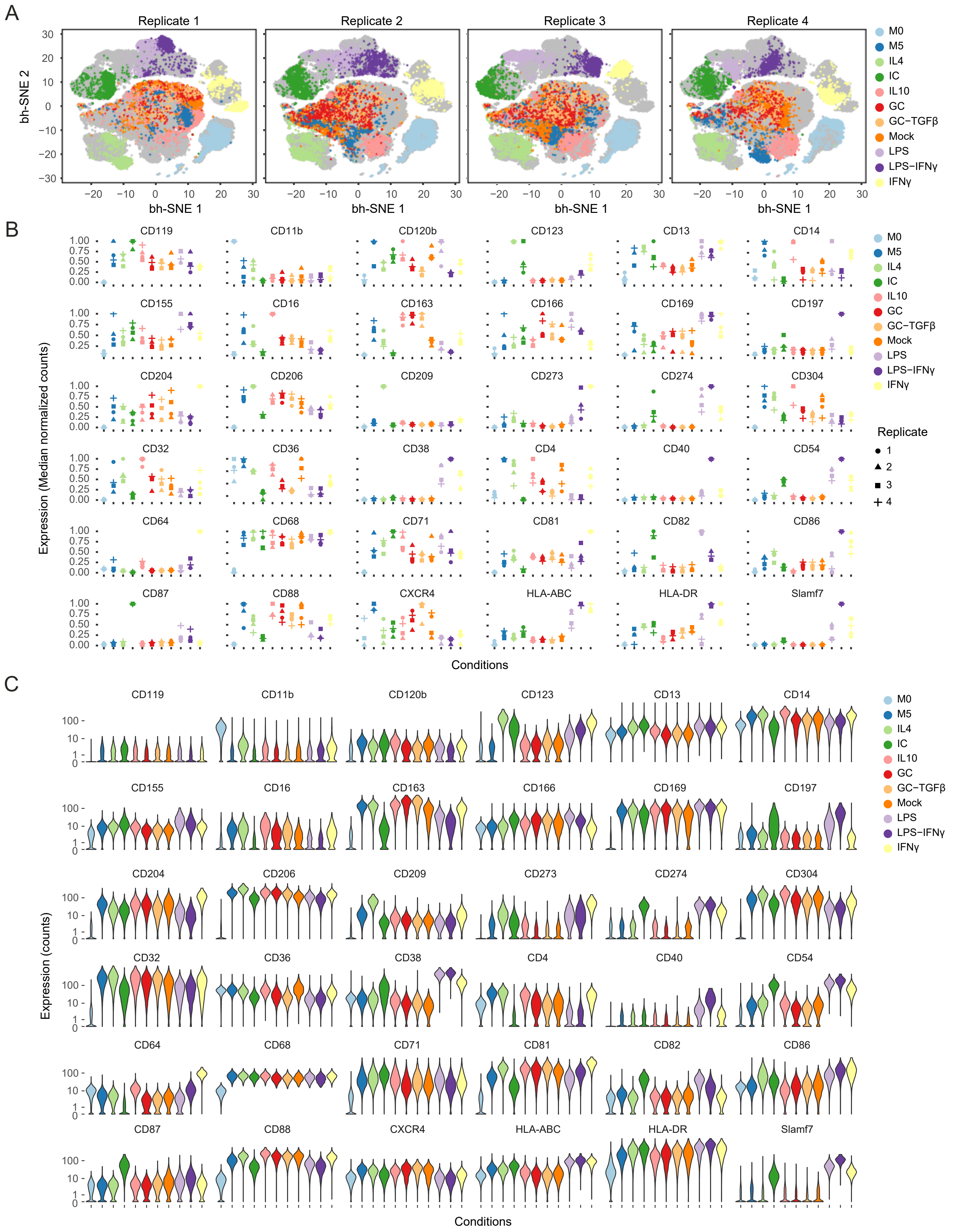


**Figure S1. Comparison of Marker Expression Across the Different MDM Populations, Related to Figure 1**

(A) The data on MDMs generated from monocytes from four healthy donors were analyzed simultaneously using the t-SNE statistic. For each sample, the data are displayed individually on a scatter plot. In each graph, the indicated replicate is displayed using the indicated color code, and the three other replicates are shown in grey. (B) Median expression levels of the indicated markers in monocytes and MDM populations across four replicates. (C) Violin plots showing the distribution of the indicated markers in the different MDM populations as assessed by mass cytometry. Data were normalized to the 99 percentile. Data shown are from one representative sample.
