## Supplementary material for "*In vitro* and *in vivo* derived macrophages occupy distinct phenotypic states": Figure S2

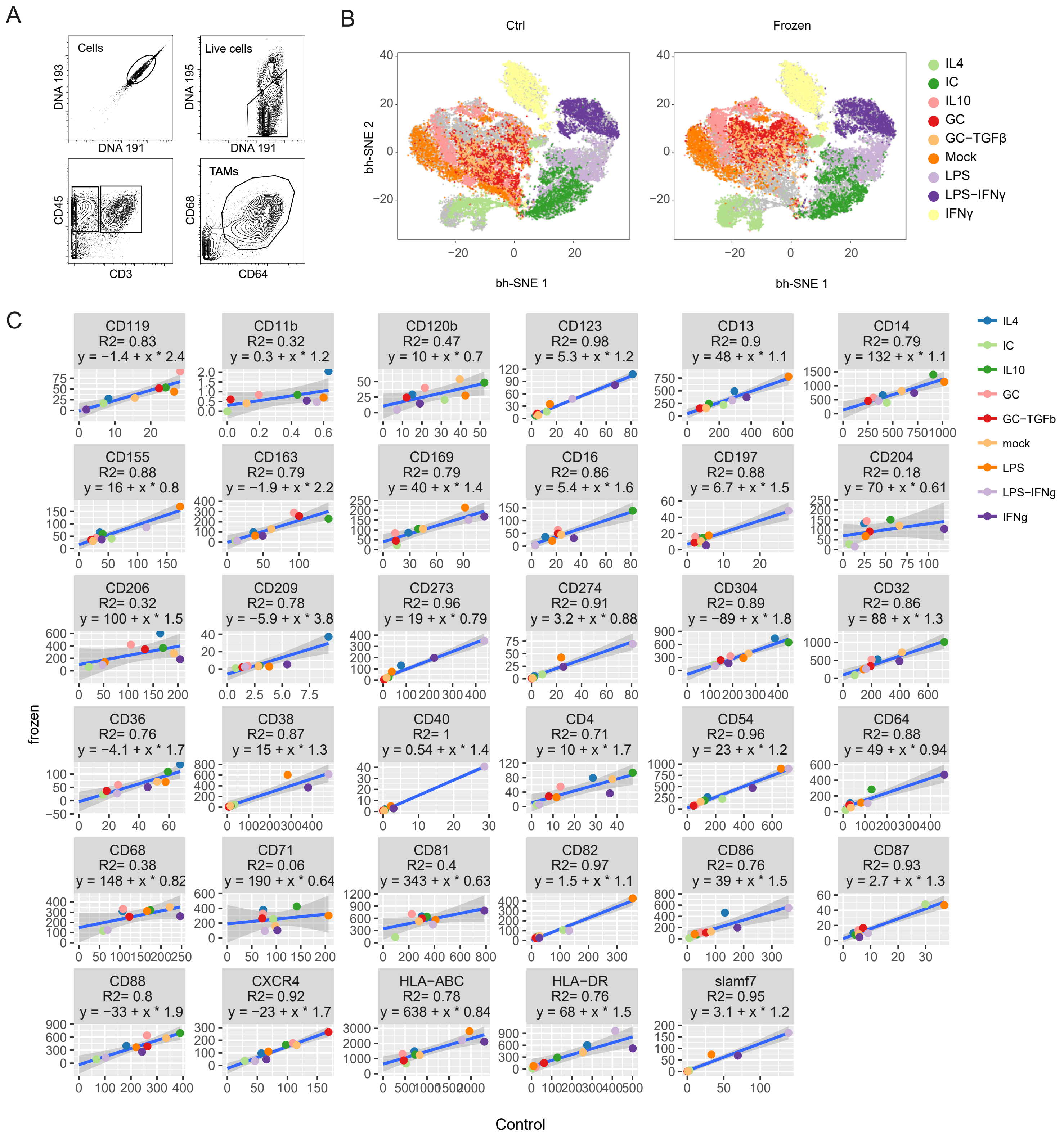


**Figure S2. Effect of Cell Freezing on Marker Expression as Assessed by Mass Cytometry, Related to Figure 2**

(A) Gating strategy based on Ir, Pt, CD45, CD3, CD64, and CD68 used to identify T cells and TAMs. (B) t-SNE maps of the different MDM populations based on the CyTOF antibody panel in presence (right panel) or in absence (left panel) of freezing step. Cells were colored per MDM sample types and in each case, the counterpart populations are colored in grey. (C) Correlation between marker expression in presence or in absence of enzymatic digestion in the indicated cell samples. For each marker, the coefficient of determination R2 and the linear model are displayed.
